# Inclusion of a 3D-printed Hyperelastic bone mesh improves mechanical and osteogenic performance of a mineralized collagen scaffold

**DOI:** 10.1101/2020.06.26.171835

**Authors:** Marley J. Dewey, Andrey V. Nosatov, Kiran Subedi, Ramille Shah, Adam Jakus, Brendan A.C. Harley

## Abstract

Regenerative repair of craniomaxillofacial bone injuries is challenging due to both the large size and irregular shape of many defects. Mineralized collagen scaffolds have previously been shown to be a promising biomaterial implant to accelerate craniofacial bone regeneration *in vivo*. Here we describe inclusion of a 3D-printed polymer or ceramic-based mesh into a mineralized collagen scaffold to improve mechanical and biological activity. Mineralized collagen scaffolds were reinforced with 3D-printed Fluffy-PLG (ultraporous polylactide-co-glycolide co-polymer) or Hyperelastic Bone (90wt% calcium phosphate in PLG) meshes. We show degradation byproducts and acidic release from the printed structures have limited negative impact on the viability of mesenchymal stem cells. Further, inclusion of a mesh formed from Hyperelastic Bone generates a reinforced composite with significantly improved mechanical performance (elastic modulus, push-out strength). Composites formed from the mineralized collagen scaffold and either Hyperelastic Bone or Fluffy-PLG reinforcement both supported human bone-marrow derived mesenchymal stem cell osteogenesis and new bone formation. Strikingly, composites reinforced with Hyperelastic Bone mesh elicited significantly increased secretion of osteoprotegerin, a soluble glycoprotein and endogenous inhibitor of osteoclast activity. These results suggest that architectured meshes can be integrated into collagen scaffolds to boost mechanical performance and actively instruct cell processes that aid osteogenicity; specifically, secretion of a factor crucial to inhibiting osteoclast-mediated bone resorption. Future work will focus on further adapting the polymer mesh architecture to confer improved shape-fitting capacity as well as to investigate the role of polymer reinforcement on MSC-osteoclast interactions as a means to increase regenerative potential.

## 1. Introduction

There is a clinical need to improve surgical repair of craniomaxillofacial bone defects. Osseous defects of the skull occur secondary to trauma, congenital abnormalities, or after resection to treat stroke, cerebral aneurysms, or cancer [1-6]. CMF defects can occur in all age ranges; cleft palate birth defects, trauma from battlefield injuries, and multiple missing teeth or oral cancer which can cause loss of jaw bone [7]. Battlefield injuries are a special class of CMF injuries, with greater than 25% of all survivable battlefield injuries in recent military conflicts in southwest Asia (Operation Iraqi Freedom, Operation Enduring Freedom) classified as maxillofacial or neck trauma, with greater than 50% attributed to explosives [8, 9]. These types of injuries result in poor healing outcomes due to infection, inadequate healing of the defect, and fixation failures [10]. Current standard of care for these defects is cranioplasty, or calvarial reconstruction, which prioritize cerebral protection over regeneration [11, 12]. Cranioplasties are common (>35,000/yr in the US; incl. >10,000 cleft palate repairs) [13, 14], but current clinical materials have significant shortcomings. Autologous or allogenic bone remains a gold standard [15, 16], but are limited by access to autologous bone [17, 18], donor site morbidity, surgical complications after cancellous autografts or cadaveric allografts (10-40%) [19, 20], and difficulty fitting irregular defects [15, 21], as well as inconsistent repair [21, 22]. Alloplastic materials are plagued by complications such as extrusion, high cost, and high infection rates (5-12x more complications than autologous transplant) [2, 3, 23-25]. These drawbacks motivate tissue engineering solutions to potentiate calvarial bone regeneration, notably metal, ceramic, and polymer-based scaffolds. Significant challenges remain, notably optimization of biomaterial strength, osteogenic activity, and ability to fit complex defect geometries. Mechanical stability and the ability to limit micromotion at the host-implant interface is crucial to the healing outcome and can directly affect osseointegration and bone regeneration [26].

Mineralized collagen scaffolds have been developed as a biomaterial implant to promote *in vivo* bone formation and *in vitro* mineral formation [27-31]. Recent work by our own laboratory has identified a mineralized collagen scaffold variant that does not need additional osteogenic supplements such as osteogenic media or Bone Morphogenic Protein-2 (BMP-2) in order to differentiate mesenchymal stem cells (MSCs) towards the osteoblastic lineage [32, 33]. These osteoprogenitors and their progeny produce a secretome to accelerate osteogenic specification, promote vascular remodeling, and suppress inflammatory damage [34], making them highly translational. We recently showed that MSC-osteoprogenitors seeded in this mineralized collagen scaffold secrete osteoprotegerin (OPG) a soluble glycoprotein and endogenous inhibitor of osteoclastogenesis and osteoclast activity; further, osteoclasts show reduced activity in response to the osteoprogenitor seeded scaffold [35]. These results suggest this mineralized scaffold may both increase MSC-osteogenesis and inhibit osteoclast activity [36]. However, despite these biological advantages the high-porosity of these scaffolds renders them mechanically weak. Successful clinical use requires the biomaterial also be surgically practical. Specifically that they be readily customized to fit complex three-dimensional defects and strong enough to withstand physiological loading [15, 37].

Beyond general mechanical strength, poor conformal contact between the biomaterial and wound margin significantly inhibits cell recruitment, angiogenesis, regenerative healing and greatly increases the risk of graft resorption [19, 38]. Implants with improved conformal contact to the host bone can limit micromotion and may improve osseointegration and regenerative healing [26]. This can be accomplished by developing shape-fitting implants that can conformally fit to the host defect site. Approaches to aid shape-fitting include the use of temperature sensitive polymers that can be shaped intraoperatively to fit complex defect sites [39-41]. Recently, our laboratory has looked to adapt three-dimensional printing approaches to create biomaterial composites with improved mechanical strength and shape-fitting capacity. Notably, we embed a mechanically-robust polymer mesh with millimeter-scale porosity into the mineralized collagen scaffold with micron-scale porosity [42]. We generated a first generation polycaprolactone (PCL) structure to form a PCL-collagen composite; this composite accelerated sub-critical defect repair in a porcine mandible defect [43, 44]. However, these PCL cages were mechanically rigid, and had no design elements to improve conformal fitting. We recently reported a 3D-printed poly(lactic-acid) based fiber with reduced percent polymer content and modifications to the fiber architecture to improve conformal fitting [45]. However, both approaches considered the polymer reinforcement phase of the composite as a passive reinforcement design rather than an active component of the biological response.

Here, we describe *in vitro* characterization of a new class of composite biomaterials. We define our composite as a mineralized collagen scaffold reinforced with a microporous mesh 3D-printed from Hyperelastic Bone or Fluffy-PLG (Dimension Inx, LLC). Fluffy-PLG is comprised of medical-grade poly(L-lactide-co-glycolide), with ≥95% internal porosity and elastic properties (2.7 ± 0.8 MPa Elastic Modulus) sufficient to form a self-supporting mesh [46], and is capable of supporting cell proliferation *in vitro* and vascularization *in vivo* [46, 47]. Hyperelastic Bone is comprised of 90 wt% calcium phosphate and 10 wt% poly(lactide-co-glycolide); Hyperelastic Bone 3D-prints have separately been shown to induce MSC osteogenic differentiation *in vitro* and new bone formation *in vivo* [48-51]. We investigate whether inclusion of 3D-printed architectures formed form Fluffy-PLG or Hyperelastic Bone 3D-paints into a mineralized collagen scaffold to create reinforced composites improves mechanical performance (stiffness, conformal fitting capacity). We subsequently examined whether inclusion of a Fluffy-PLG or Hyperelastic Bone mesh provides biological advantage via degradation by-products to improve osteogenic response of human mesenchymal stem cells (hMSCs) within the mineralized collagen scaffold. These studies therefore consider inclusion of architectured cellular structures into mineralized collagen scaffolds to provide significant mechanical and biological advantage for regenerative medicine applications.

## 2. Materials and Methods

### 2.1. Experimental Design

Composites were fabricated from mineralized collagen scaffolds combined with reinforcing support architectures 3D-printed from Hyperelastic Bone or Fluffy-PLG 3D-paints (Dimension Inx LLC, Illinois, USA) (**Fig. 1A**). Throughout this study we compared the three material groups: mineralized collagen scaffolds (MC) on their own, mineralized collagen composites reinforced with 3D-printed Hyperelastic Bone, and mineralized collagen composites reinforced with 3D-printed Fluffy-PLG (MC-Fluffy). Although nano-and microstructurally distinct, 3D-printed Fluffy-PLG and Hyperelastic Bone are based on the same polylactide-co-glycolide (PLG), with the only compositional difference being addition of 90wt% calcium phosphate mineral in the Hyperelastic Bone. Studies of the effect of degradation products (released lactic and glycolic acid; changes in pH; calcium and phosphorous release) or degradation induced changes in mechanical performance (elastic modulus) was evaluated for 3D-printed Hyperelastic Bone. *In vitro* testing of hMSC osteogenesis and mineral formation was performed using mineralized collagen scaffolds as a function of the inclusion of 3D-printed, reinforcing Hyperelastic Bone or Fluffy-PLG the form of a cross-design (**Fig. 1B**). Overall mechanical performance (elastic modulus; shape-fitting ability) was performed on reinforced composites with a 3D-printed mesh design, and was evaluated via compression and push-out tests to determine if inclusion of the reinforcing structures improve mechanical properties of the mineralized collagen scaffolds (**Fig. 1B**).

**Fig. 1.**
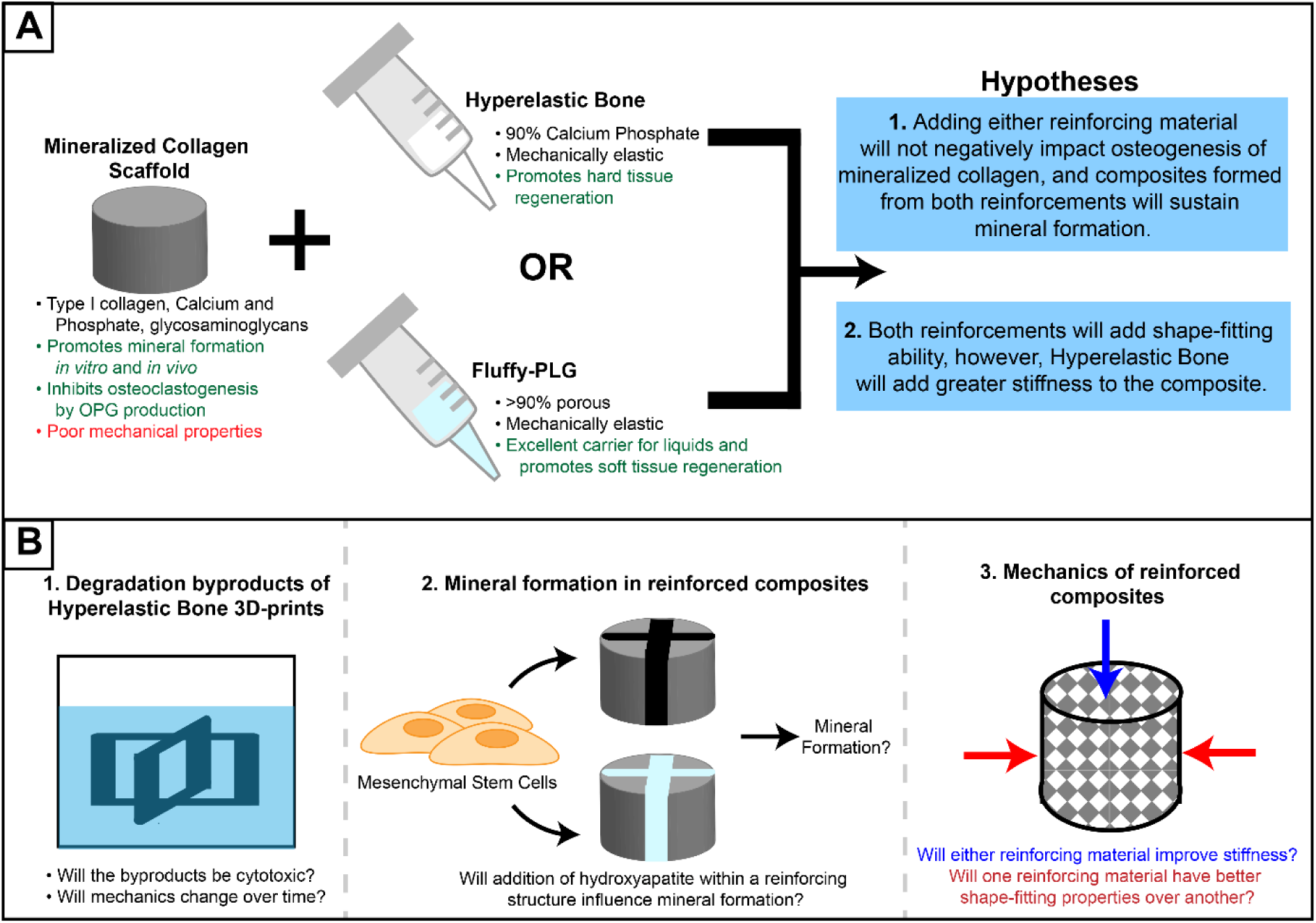
Experimental outline. Mineralized collagen scaffolds promote bone repair, however, these have poor mechanical properties including lack of shape-fitting behavior. (A) We examined incorporating reinforcing structures 3D-printed from either Hyperelastic Bone or Fluffy-PLG, to the mineralized collagen scaffolds. We investigated (1) if addition of the reinforcing structures could increase the osteogenic response, and if the 3D-printed structure containing calcium phosphate would promote bone formation over the one without and (2) the influence of the 3D-printed structures on scaffold mechanical properties (i.e. compressive properties and shape-fitting). (B) Steps of the experiment to answer the study questions.

### 2.2. 3D-printing Hyperelastic Bone and Fluffy-PLG fiber arrays

Hyperelastic Bone and Fluffy-PLG 3D-Paints were provided by Dimension Inx LLC and 3D-printed using a Manufacturing Series 3D-BioPlotter (EnvisionTEC, Michigan, USA). Constructs used for degradation studies and *in vitro* culture were printed as cross designs, symmetrical crosses 6 mm in diameter and 3 mm in height with a 0.7 mm feature thickness (**Fig. 2**) using a 27 Ga nozzle at a speed of 2-5 mm/s dependent on solution viscosity, a room temperature stage and deposition, and low print speeds to accommodate the fine structure of the print [46, 47, 51]. These structures display low polymer volume fractions within the resulting composite (13.61 v/v%). Hyperelastic Bone 3D-prints were stored in the dark at 4°C until use while Fluffy-PLG 3D-prints were stored at −80°C until use, per manufacturer instructions. A 3D-printed mesh design was used to reinforcement mineralized collagen scaffolds for mechanical testing. These were prepared and printed by Dimension Inx LLC as square sheets at 70 mm/s (60 mm on a side; 6 mm thick), with a standardized 120° angle [47]. 3D-printed reinforcing meshes were printed with Fluffy-PLG and prepared with two distinct feature sizes (2 mm or 3 mm line spacing). However, Hyperelastic Bone 3D-printed meshes were only tested for 2 mm line spacing (3 mm line spacing composites were not stable with regard to handling and manipulation due to the large structural pore size relative to the small, comprising fiber diameters). A 2 mm line spacing print is approximately 83% porous based on the design, not including inherent material porosity.

**Fig. 2.**
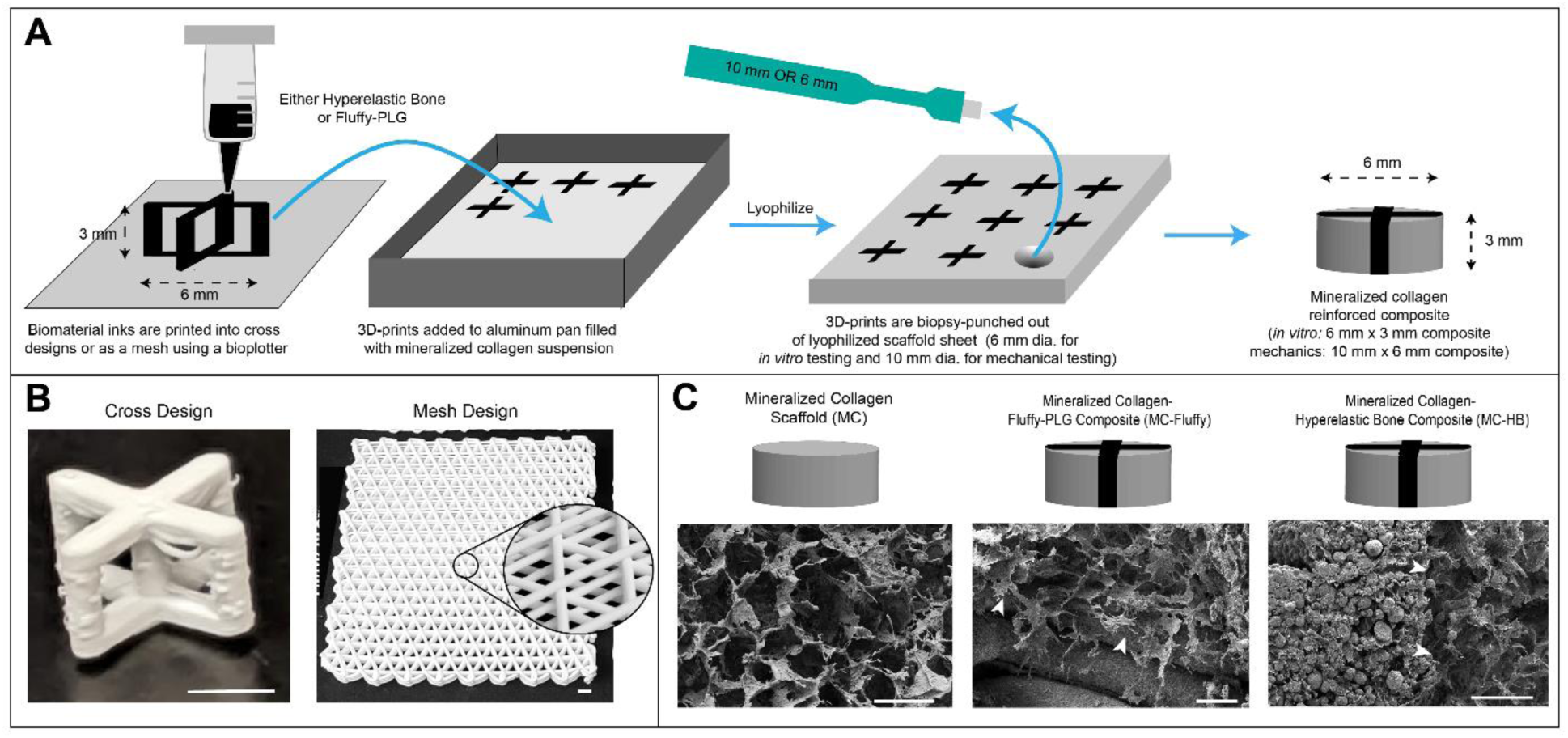
Fabrication of composites used *in vitro* and for mechanical testing. (A) Hyperelastic Bone and Fluffy-PLG were 3D printed in “cross designs” for *in vitro* testing or as a mesh for mechanical testing. Cross prints were printed individually and added to a continuous mineralized collagen suspension in an aluminum mold. After the addition of multiple prints, the entire mold was freeze-dried and a 6 mm biopsy punch was used to remove scaffold and 3D-print together. Mesh reinforced composites were fabricated by adding the entire mesh to the aluminum mold with mineralized collagen suspension and biopsy punching 10 mm diameter composites after lyophilization. Mesh 3D-prints were printed with either 2 mm or 3 mm line spacing. (B) Images of cross design and mesh 3D-prints, scale bar represents 3 mm. (C) SEM images of mineralized collagen scaffold and cross design reinforced composites and acronyms used to represent the groups in the study. Scale bar represents 150 µm and white arrows indicate interface between the mineralized collagen and 3D-printed reinforcing structures.

### 2.3 Analysis of degradation byproducts and changes in mechanical properties of Hyperelastic Bone 3D-prints

Degradation byproducts of 3D-printed Hyperelastic Bone were analyzed using a standardized ‘cross design’ print (**Fig. 2**). Prints were washed twice in 70% ethanol for 30 min and then twice in cell grade water for 30 min per the manufacturer’s instructions before submerging individual mesh structures in 5 mL of Phosphate Buffered Saline (PBS) in individual glass vials. Sample were then either maintained at 37°C in an incubator (standard degradation; to simulate degradation during *in vitro* cell culture or *in vivo*), or in a 90°C oven (Isotemp Vacuum Oven Model 282A, Fisher Scientific, Massachusetts, USA) to facilitate accelerated degradation [52].

#### pH

The pH of 3D-print conditioned PBS samples from the 90°C degradation study was measured using a FiveEasy Standard pH meter (Mettler Toledo, Ohio, USA) for n = 8 samples (PBS as a control).

#### Lactic and glycolic acid release

Over the course of a 28-day accelerated release experiment (‘cross design’ prints in 5 mL PBS at 90°C), the PBS incubation solution was centrifuged at regular intervals to isolate the liquid supernatant for analysis of lactic and glycolic acid concentrations. 5 mL of PBS, along with any particles separated during centrifugation, were added to each sample vial until the next isolation point. Colorimetric determination of lactic acid concentration followed procedures in literature [53]. Briefly, a standard curve was created using a solution of 1% L-(+)-Lactic Acid (Sigma Aldrich, Missouri, USA) and half-dilutions. PBS and iron (III) chloride (Sigma Aldrich) were used as controls. 50 µl of sample was mixed with 2 mL of 0.2% iron (III) chloride, and absorbance was measured at a wavelength of 390 nm on spectrophotometer (Tecan, Switzerland). Colorimetric determination of glycolic acid release followed procedures in literature [54]. A 1 mg/mL solution of glycolic acid (Sigma Aldrich) was used to create half-dilutions to generate a standard curve. 500 µL of sample was evaporated to dryness at 125°C in a vacuum oven at atmospheric pressure. β-Naphthol (Sigma Aldrich) in 92% sulfuric acid was added to vials containing evaporated sample and boiled for 20 min. After boiling, 80% sulfuric acid was added to samples for 10 min before measuring absorbance at a wavelength of 480 nm on a spectrophotometer (Tecan). A sample size of n = 8 was used for both lactic acid and glycolic acid release measurements.

#### Calcium and phosphate release

Calcium and Phosphate release from 3D-printed Hyperelastic Bone and mineralized collagen scaffolds were quantified through a 28 days standard release experiment (culture in 5 mL of PBS at 37°C). 1 mL PBS aliquots were removed from suspension, then combined with 2 mL of Trace Metal Grade concentrated HNO_3_ (Thermo Fisher Scientific 67-70%) for a 30-minute pre-digestion step. The tubes were then capped and placed into a rotating carousel inside MARS 6 (CEM Microwave Technology Ltd., North Carolina, USA) microwave digester (40 minutes). The final digested solution was diluted to a volume of 50 mL in DI water, then analyzed via inductively coupled plasma-mass spectrometer (ICP-OES, Optima 8300, PerkinElmer, USA; elemental analysis in axial mode; **Supp. Tables 1 and 2**). Eight samples of each group were used per timepoint.

#### Mechanical analysis

Mechanical testing was performed on Hyperelastic Bone 3D-prints after 0, 7, and 28 days in PBS at 37°C (PBS was changed every 3 days). Prints were aerated in the incubator overnight and then dried at room temperature overnight before testing. Compression testing was performed with an Instron 5943 mechanical tester (Instron, Massachusetts, USA) using a 100 N load cell at a rate of 1 mm/min. Seven samples at each timepoint were tested and stress-strain curves were analyzed using a custom Matlab program to determine Young’s modulus, ultimate stress, and ultimate strain.

### 2.4. Fabrication of mineralized collagen scaffolds and reinforced composites

Mineralized collagen scaffolds were fabricated via lyophilization from a liquid suspension as previously described [27, 28, 32, 55, 56]; while composites were fabricated via lyophilization as previously described [43-45]. Briefly, 1.9 w/v% type I collagen from bovine Achilles tendon (Sigma Aldrich) was blended together with a 40 wt% mineral suspension of phosphoric acid (Fisher Scientific) and calcium hydroxide (Sigma Aldrich), 0.84 v% chondroitin-6-sulfate sodium salt from shark cartilage (Sigma Aldrich), and additional calcium nitrate tetrahydrate (Sigma Aldrich) using a rotor-stator in a jacketed cooling vessel until smooth [27, 28, 42, 43]. Mineralized collagen scaffolds for *in vitro* and mechanical testing were fabricated by pipetting the suspension into aluminum pans and lyophilizing using a Genesis freeze-dryer (VirTis, New York, USA). Scaffolds were added to the freeze-dryer shelf at 20°C and the temperature was dropped at a rate of 1°C/min to −10°C in order to form ice crystals. Samples were held at −10°C for 2 hours and then ice crystals were sublimated by decreasing the pressure and temperature to create a porous scaffold. After sublimation, solid scaffolds were brought back to room temperature and atmospheric pressure before storing in a desiccator prior to use. Composites (MC-HB or MC-Fluffy) were formed by adding the 3D-printed reinforcement, either Hyperelastic

Bone or Fluffy-PLG, into the collagen suspension prior to lyophilization (**Fig. 2**). Prior to incorporation into the mineralized collagen suspension, 3D-prints were washed per supplier instructions: Hyperelastic Bone 3D-prints in 70% ethanol then cell grade water; Fluffy-PLG prints in cell grade water to remove salts, then 70% ethanol, then in cell grade water again [46, 51]. Lyophilization conditions to create composites used the same temperature and pressure profiles as scaffolds. Scaffolds and composites for *in vitro* testing were biopsy-punched out of the resulting sheet with a 6 mm diameter biopsy punch, while scaffolds and composites for mechanical testing used a 10 mm diameter biopsy punch.

### 2.5. Scanning electron microscopy of collagen scaffolds and reinforced composites

Scanning electron microscopy (SEM) was used to visualize collagen infiltration into the 3D-printed structures. Dry collagen scaffolds and reinforced composites (MC-HB, MC-Fluffy) were cut to expose the interior before sputter coating with Au/Pd (Denton Desk II TSC, New Jersey, USA). After sputter-coating, samples were imaged using an FEI Quanta FEG 450 ESEM (FEI, Hillsboro, OR). Both composites reinforced with cross design and mesh design 3D-prints were imaged for collagen infiltration into 3D-prints (**Fig. 2, Supp. Fig. 2**).

### 2.6. Compression and shape-fitting testing of collagen scaffolds and reinforced composites

Mechanical performance of collagen reinforced with a standardized 120° mesh morphology 3D-print was evaluated [47]. Scaffolds and reinforced composites formed from either Hyperelastic Bone or Fluffy-PLG were tested under mechanical compression to determine elastic modulus as well as using a standardized push-out test. Reinforced composites and scaffolds measured approximately 10 mm in diameter by 6 mm in height, and both mechanical tests used an Instron 5943 mechanical tester (Instron) with a 100 N load cell. Mechanical compression was performed at a rate of 1 mm/min and the linear portion of stress-strain curves was analyzed using a custom Matlab program to determine Young’s Modulus [43, 45]. Push out testing was performed at a rate of 2 mm/min following previously described methods [45, 57]. Briefly, samples were compressed and fit into an 8.5 mm diameter hole in a Teflon plate and samples were pushed through the mold to measure maximum load achieved to move the samples. Eight samples were used for each test in order to determine stiffness and shape-fitting ability.

### 2.7. Cell culture and biomaterial preparation

Bone marrow derived human mesenchymal stem cells (BM-hMSCs) (24yr old B female, Lonza, Switzerland) were used at passage 4 – 6 and cultured with complete hMSC media containing low glucose Dulbecco’s Modified Eagle Medium and glutamine (School of Chemical Sciences Cell Media Facility, University of Illinois), Fetal Bovine Serum (Gemini Bio Products, California, USA), and antibiotic-antimycotic solution (Thermo Fisher Scientific, Massachusetts, USA). Cell contamination was tested with a MycoAlert™ Mycoplasma Detection Kit (Lonza) and cells tested negative for mycoplasma.

Scaffolds and reinforced composites were sterilized via ethylene oxide treatment with an AN74i Anprolene gas sterilizer (Andersen Sterilizers Inc., North Carolina, USA) for *in vitro* testing. Prior to adding cells to scaffolds and reinforced composites, these followed a standard hydration procedure for mineralized collagen scaffolds previously reported [45, 58-60]. Briefly, samples were hydrated in 70% ethanol, washed in PBS, crosslinked with 1-Ethyl-3-(3-dimethylaminopropyl) carbodiimide and N-Hydroxysuccinimide, followed by washing in PBS, and finally soaking in hMSC complete media for 2 days.

### 2.8. Influence of Hyperelastic Bone degradation byproducts on cell activity

Cytotoxic effects of Hyperelastic Bone degradation byproducts were determined via a previously defined elution assay [61]. Briefly, Hyperelastic Bone structures were printed into standard ‘cross-designs’ (**Fig. 2**) then placed in PBS either in a 37°C incubator or in a 90°C vacuum oven (accelerated degradation) for 28 days to create degradation byproduct conditioned media. Cell cytotoxicity was determined using for 10,000 MSCs in individual wells of a 96-well plate. Briefly, hMSCs were first cultured in complete hMSC media for 24 hrs. After 24 hrs, media was removed and replaced for 24 hours with PBS containing eluted degradation factors at full strength (100%) or conditioned PBS diluted with hMSC media (50% sample, 25% sample, and 12.5% sample). Non-conditioned PBS was used as a control. After 24 hours, metabolic health of hMSCs was examined via an alamarBlue™ (Thermo Fisher Scientific) assay, measuring the fluorescence of resorufin (540 nm excitation, 580 nm emission) using a F200 spectrophotometer (Tecan). Metabolic activity was compared to a standard curve of known cell numbers. There were 8 samples of each group tested with PBS used as a control.

### 2.9. Cell activity within scaffolds and reinforced composites

Scaffolds and reinforced composites were added to 24-well plates and seeded via a standard static seeding assay (5,000 cells/µL per side; 100,000 cells/scaffold) using previously defined procedures [45, 58]. After allowing cell attachment, hMSC complete media was added to wells and plates containing cell-seeded scaffolds and then were added to a 37°C incubator. The metabolic activity of cell-seeded biomaterials was examined via an alamarBlue™ (Thermo Fisher Scientific) assay at days 1, 4, 7, 14, and 28. Briefly, scaffolds and composites were washed in PBS, followed by a 1.5 hr soak in alamarBlue™ and complete hMSC cell media on a shaker in a 37°C incubator [45, 58]. A standard curve of known cell number was used to calculate fold change of metabolic activity over the initial cell seeding density (a fold change of 1 represented the metabolic activity of 100,000 cells). Six samples were used to determine metabolic activity, and these were used for the entirety of the study (non-destructive assay).

Expression levels of cell secreted Osteoprotegerin and Vascular Endothelial Growth Factor (VEGF) was examined via ELISA (R&D Systems, Minnesota, USA). Briefly, media was collected and replaced from cell-seeded samples every 3 days until day 28 to determine cumulative protein expression. ELISAs were performed to using 25 µL of sample and 75 µL reagent diluent and compared against a known standard curve to quantify expression level. Six samples were used throughout the study (non-destructive).

Analysis of gene expression profiles was performed at days 1, 4, 7, 14, and 28 of culture. Specimens were washed in PBS, frozen at −80°C, pulverized on dry ice with disposable pestles (Thermo Fisher Scientific), then treated with an RNeasy Plant Mini Kit (Qiagen, California, USA) [62]. Concentrations of isolated RNA were measured using a Nanodrop Lite (Thermo Fisher Scientific). Reverse transcription of RNA to cDNA was performed following directions and supplies from a QuantiNova Reverse Transcription kit (Qiagen) and a S100 thermal cycler (Bio-Rad, Hercules, California). After reverse transcription, PCR was performed on cDNA to quantify gene expression. 10 ng of cDNA was used in each well and Taqman primers were purchased from Thermo Fisher Scientific (*RUNX2, COL1A2, Osterix, FGFR2, IGF2*) with *GAPDH* serving as a housekeeping control (**Supp. Table 3**). Plate preparation was performed using a Gilson Pipetmax liquid handling machine (Gilson, Wisconsin, USA) and plates were read using a QuantstudioTM 7 Flex Real-Time PCR System (Thermo Fisher Scientific). Data was analyzed using the delta-delta CT method to generate box plots for a fold change of gene expression (with a fold change of 1 representing the gene expression of 100,000 hMSCs before seeding on scaffolds and composites). Five samples were used at each timepoint.

### 2.10. Analysis of mineral formation in scaffolds and reinforced composites

Inductively Coupled Plasma (ICP) Optical Emission spectrometry was performed to assess mineral formation at the end of *in vitro* culture (day 28). MC, MC-HB, and MC-Fluffy samples were washed in PBS, fixed with formalin (Formal-Fix, Thermo Fisher Scientific) for 24 hrs at 4°C, washed again in PBS, then dried on a Kimwipe before storing at −80°C until use. Before performing ICP, samples were lyophilized at the same conditions used to fabricate the original scaffolds. Samples were weighed then dissolved using concentrated nitric acid, Trace Metal Grade concentrated HNO_3_ (Thermo Fischer Scientific 67-70%), followed by automated sequential microwave digestion in a CEM Mars 6 microwave digester. The acidic solution was diluted to a volume of 50 mL using DI water, so as to make the final concentration of the acid <5%. The ICP-OES was calibrated with a series of matrix matched standards before introducing the unknown samples. Digestion and ICP-OES analysis parameters are listed in **Supp. Tables 4 and 5**. Nine samples were used for each group and these were normalized to the calcium and phosphorous content of respective dry scaffolds and composites without cells in order to get a fold change and new calcium and phosphorous deposition.

### 2.11. Statistical Methods

Statistics followed procedures outlined by Ott and Longnecker [63]. Quantity of samples used was based off previous experiments using similar sample groups and a 95% confidence interval for all tests [45, 58]. For all data, normality was evaluated and if data was not normal a Grubb’s outlier test was performed and normality was re-assessed. Analysis of more than two groups used an ANOVA, and depending on whether assumptions of normality (Shapiro-Wilk) and equal variance (Levene’s Test) of residuals was met, a specific ANOVA was used as outlined in **Supp. Table 6**. For data involving two groups, either a paired T-test or a two-sample T-test was used. Normality (Shapiro-Wilk) and two sample T-test for variance were completed using OriginPro software (OriginLab, Massachusetts, USA) before analysis. For non-normal data a paired sample Wilcoxon Signed Rank test was used or a two-sample Kolmogorov-Smirnov test was used. For samples with normal data and unequal variance, a two sample T-test with a Welch correction was used as outline in **Supp. Table 7**. For powers lower than 0.8, data was deemed inconclusive. Data is expressed as average ± standard deviation unless otherwise noted.

## 3. Results

### 3.1. Degradation byproducts of Hyperelastic Bone does not negatively affect cell viability

Hyperelastic Bone 3D-printed as a ‘cross design’ completely degraded over the course of a 28 day-accelerated degradation study (PBS; 90°C). Aliquots taken from the Hyperelastic Bone degradation experiments were first analyzed to define changes in pH as well as glycolic acid and lactic acid elution; changes in pH were also compared to a PBS standard (pH 7.4) as well as mineralized collagen scaffolds exposed to accelerated degradation conditions (PBS; 90°C). Overall, both mineralized collagen scaffolds and Hyperelastic Bone 3D-prints drove a drop in solution pH during early stages of degradation; scaffolds showed a sharper decrease in pH while the Hyperelastic Bone showed a temporally extended drop in pH. However, acidic byproducts were only detectable during the first week of degradation (**Supp. Fig. 1A**). Analysis of lactic and glycolic acid content in the media suggested the majority released lactic and glycolic acid occurred rapidly as well, with a total of 12.7 mg lactic acid and 307.5 µg of glycolic acid released (**Supp. Fig. 1B, C**). Finally, we investigated whether the degradation byproducts from the Hyperelastic Bone structures drove a measurable change in metabolic activity of hMSCs. The total eluted byproducts from Hyperelastic bone structures in PBS were collected from both standard (37°C) and accelerated (90°C) degradation protocols over 28 days and compared to PBS. hMSCs were cultured in a mixture of conventional cell culture media and PBS (PBS; 37°C degradation byproducts; 90°C degradation byproducts) at discrete ratios: 12.5% PBS, 25% PBS, 50% PBS, 100% PBS. While hMSC metabolic activity reduced with increasing amounts of PBS (vs. cell culture media), there was no significant trend suggesting a decrease in hMSC activity as a function of Hyperelastic Bone degradation byproducts (**Supp. Fig. 1D**).

### 3.2. 3D-printed Hyperelastic Bone 3D-prints lose stiffness over time and release less calcium than mineralized collagen scaffolds

We subsequently quantified degradation-induced changes in compressive properties and release of calcium and phosphorous ions from Hyperelastic Bone 3D-printed as a cross design over the course of a 28-day standard degradation experiment (37°C, PBS). While there were non-significant decreases in Young’s modulus over the initial 7 days, significant degradation of the constructs over the full 28-day experiment were marked by significant decreases in Young’s modulus as well as average ultimate stress and strain (**Table 1**).

**Table 1.** Youngs Modulus, ultimate stress, and ultimate strain of Hyperelastic Bone 3D-printed cross designs in PBS at 37°C for 7 and 28 days compared to prints not soaked. * denotes that day 28 prints had irregular stress-strain curves and multiple prints snapped before compression testing. Day 0 Young’s Moduli and was significantly (p < 0.05) different from Day 28 Young’s Moduli. Day 0 Ultimate Stress and Ultimate Strain were significantly (p < 0.05) greater than the day 7 and day 28 groups. Data expressed as mean ± standard deviation (n=7).

|  | Average Young's Modulus (MPa) | Average Ultimate Stress (MPa) | Average Ultimate Strain (mm/mm) |
| --- | --- | --- | --- |
| **Day 0** | 4.8 ± 1.3 | 0.76 ± 0.27 | 0.27 ± 0.03 |
| **Day 7** | 3.1 ± 1.4 | 0.32 ± 0.16 | 0.18 ± 0.04 |
| **Day 28** | 1.3 ± 1.0 * | 0.09 ± 0.03 | 0.13 ± 0.04 |

Further, the structures were difficult to handle after 28 days of exposure, with multiple breaking prior to mechanical testing. Significant elution of both calcium and phosphorous was observed for Hyperelastic Bone structures. Interestingly, while significant calcium (p < 0.05) and phosphorous was released from the Hyperelastic Bone structures, overall release was less, significantly in the case of calcium, than that released from the native mineralized collagen scaffold in the same conditions (**Table 2**). However, this release suggests the embedded Hyperelastic Bone component may supplement the mineral ions released from the mineralized collagen scaffold phase to aid osteogenesis.

**Table 2.** Calcium and Phosphorous release from mineralized collagen scaffolds and Hyperelastic Bone 3D-prints at 37°C in PBS after 28 days. * indicates the calcium release in mineralized collagen scaffolds is significantly (p < 0.05) greater than the calcium release in Hyperelastic Bone 3D-prints. Data expressed as mean ± standard deviation (n=8).

| Sample | Calcium Released (ppm) | Phosphorous Released (ppm) |
| --- | --- | --- |
| Mineralized Collagen | 32.8 ± 8.5 * | 42.6 ± 11.2 |
| Hyperelastic Bone | 2.6 ± 0.7 | 25.1 ± 15.7 |

### 3.3. Integration of 3D-printed structures and lyophilized scaffold to form a composite

Reinforced collagen composites were formed using 3D-printed structures generated in either a conventional ‘Cross Design’ for *in vitro* trials or using a standardized 120° ‘Mesh Print’ (Dimension Inx) for mechanical analysis. SEM analysis of both cross design and mesh design 3D-print composites (MC-HB, MC-Fluffy) showed close integration of the mineralized collagen microstructure with the Hyperelastic Bone or Fluffy-PLG printed structure (**Fig. 2C, Supp. Fig. 2**).

### 3.4. Hyperelastic Bone composites show improved mechanical performance

Mechanical compression and push-out tests were performed on scaffolds and composites formed from a symmetrical 120° mesh morphology. MC-HB composites displayed significantly (p < 0.05) greater stiffness than MC-Fluffy composites or the native mineralized collagen scaffolds (**Fig. 3A**), while inclusion of a Fluffy-PLG 3D-print afforded no increase in modulus relative to the mineralized collagen scaffold on its own. MC-HB composites also displayed increased push-out force than MC-Fluffy composite or the scaffold (**Fig. 3B**). Altering the mesh spacing of MC-Fluffy composites had no effect on stiffness or shape-fitting ability of the reinforced composite (**Supp. Table 8**).

**Fig. 3.**
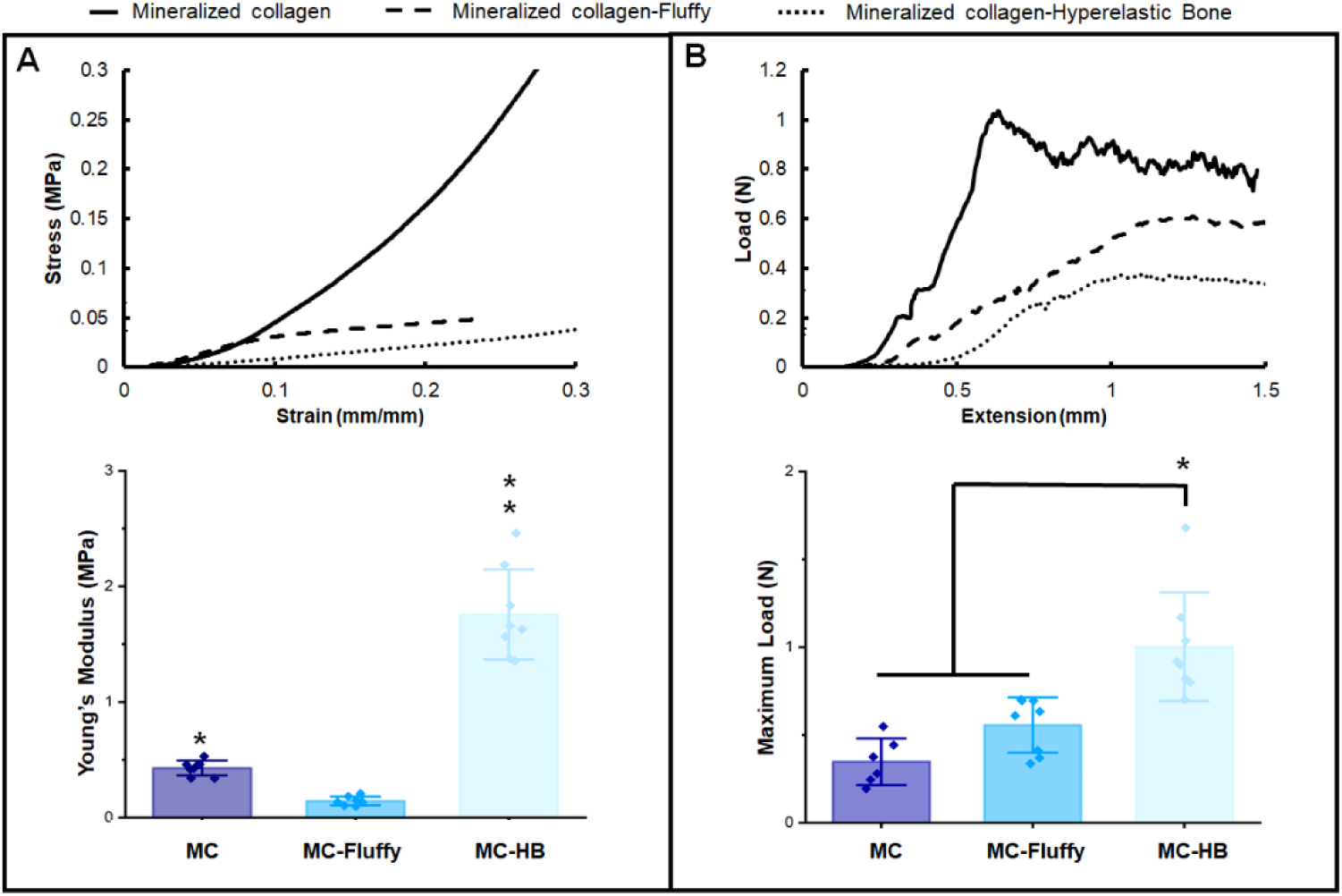
Mechanical behavior of 3D-printed mesh reinforced composites. Compression testing to determine Young’s Modulus and shape-fitting testing to determine maximum load during specimen push-out was performed on mineralized collagen (MC), Hyperelastic Bone reinforced composites (MC-HB), and Fluffy-PLG reinforced composites (MC-Fluffy). (A) Young’s Modulus averages and representative stress-strain curves from compression testing. Asterix represents all groups are significantly (p < 0.05) different from one another. (B) Average maximum load and representative load-extension curves from push-out testing. * represents the MC-HB group is significantly (p < 0.05) greater than both the MC and MC-Fluffy groups. Error bars represent average ± standard deviation (n=8).

### 3.5. Reinforced composites support hMSC metabolic activity and promote increased osteogenic activity

Both the native mineralized collagen scaffold as well as composites formed from either Fluffy-PLG or Hyperelastic Bone 3D-prints supported significant increases in the metabolic activity of seeded hMSCs over 28 days *in vitro* experiment (**Fig. 4A**). Interestingly, MC-HB and MC-Fluffy composites showed significantly (p < 0.05) greater metabolic activity relative to the mineralized collagen scaffold alone for days 4 – 14. However, by day 28 all groups (MC, MC-HB, MC-Fluffy) showed approximately 3-fold increases in metabolic activity versus the start of the experiment, with no significant differences between groups.

**Fig. 4.**
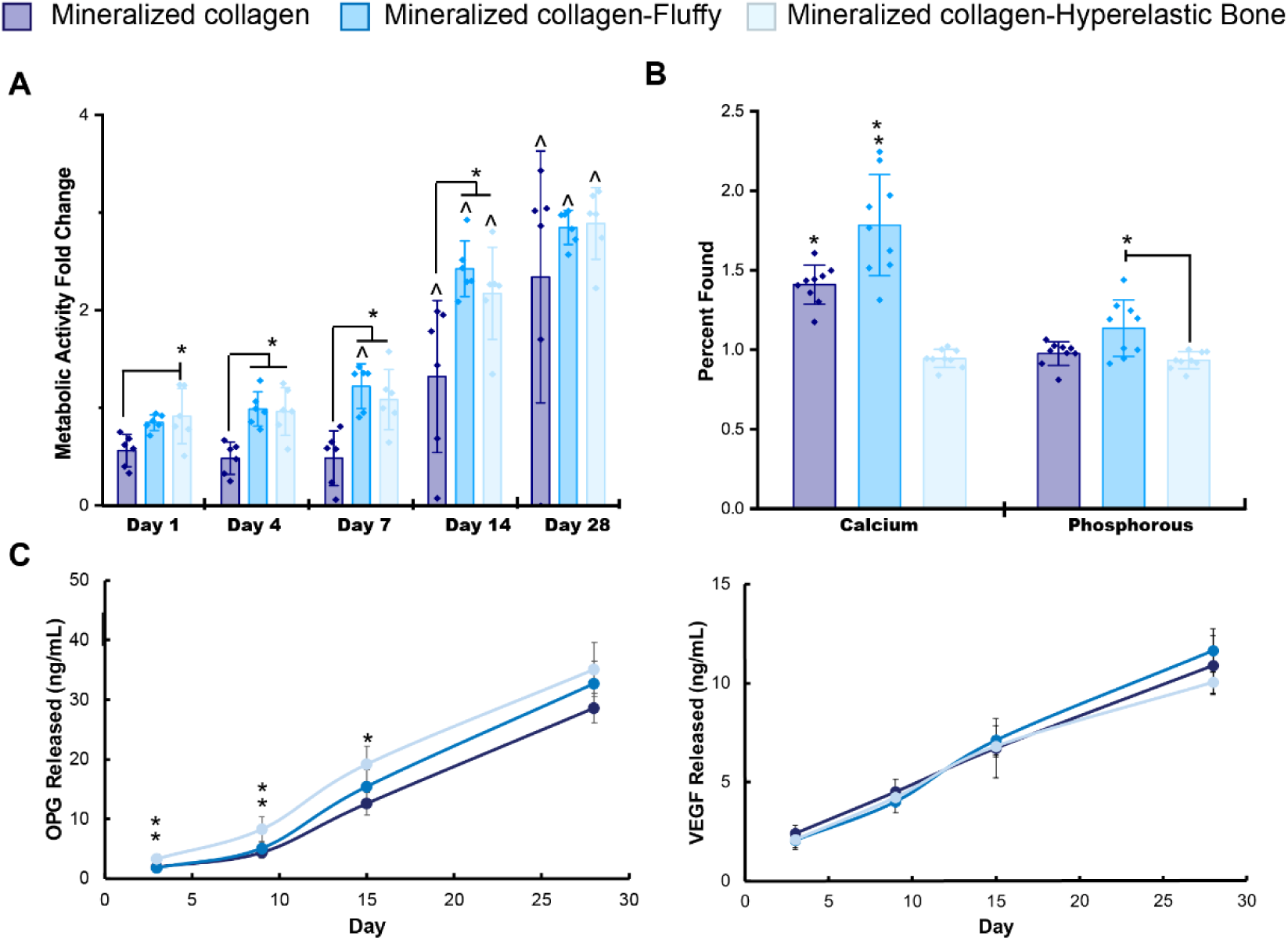
Metabolic activity, mineral formation, and protein expression of hMSCs seeded on mineralized collagen scaffolds (MC), Fluffy-PLG reinforced composites (MC-Fluffy), and Hyperelastic Bone reinforced composites (MC-HB). (A) The metabolic activity of 100,000 hMSCs is denoted by an activity of 1, with the y-axis representing a fold change in activity over this cell seeding density. * indicates one group is significantly (p < 0.05) greater than another group. ^ indicates one group is significantly (p < 0.05) greater than the same group compared to day 1. Data expressed as average ± standard deviation (n=6). (B) Percent found of Calcium (Ca) and Phosphorous (P) was determined by ICP analysis of groups seeded with cells after 28 days and normalized to the respective day 0 unseeded groups. Asterix indicate significance and all groups have significantly (p < 0.05) different calcium content. The MC-Fluffy group has significantly (p < 0.05) more phosphorous than the MC-HB group. Data expressed as average ± standard deviation (n=9). (C) Protein expression in scaffolds and composites was analyzed with an ELISA for OPG and VEGF released from pooled media over 28 days. ** indicates the MC-HB group was significantly (p < 0.05) greater than the other two groups at day 3 and 9. * indicates the MC-HB group was significantly (p< 0.05) greater than the MC group at day 15. There were no significant (p < 0.05) differences in VEGF released between all tested groups. Data expressed as average ± standard deviation (n=6).

We quantified release profiles for VEGF and OPG from the MSC-seeded scaffold or composites (**Fig. 4C**). While steady increase in VEGF released into the media was observed for all constructs, there was no significant difference between release profiles for all groups. However, we observed significantly (p < 0.05) increased OPG released from hMSC-seeded MC-HB composites versus both hMSC-seeded MC scaffolds (days 3, 9, 15) or hMSC-seeded MC-Fluffy composites (days 3 and 9). However, by day 28 the effect of the Hyperelastic Bone composite on OPG production was no longer significant.

### 3.6. Scaffolds and composites promote osteogenic gene expression

We examined temporal expression profiles for a series of genes associated with hMSC osteogenic specification: *RUNX2, Osterix, FGFR2, COL1A2*, and *IGF2* (**Fig. 5**). Expression profiles between groups were largely similar. However, MC-HB composites promoted significantly (p < 0.05) greater expression fold changes for *RUNX2* (day 1), *Osterix* and *FGFR2* (days 1, 14), and reduced expression of *COL1A2* (days 4, 14) compared to mineralized collagen scaffolds. MC-Fluffy composites promoted significantly (p < 0.05) greater expression fold changes for *RUNX2* (day 7) and reduced expression of *COL1A2* (day 14) compared to mineralized collagen scaffolds. MC-HB composites also promoted significantly (p < 0.05) greater expression of *IGF2* than MC-Fluffy composites and MC scaffolds at day 1.

**Fig. 5.**
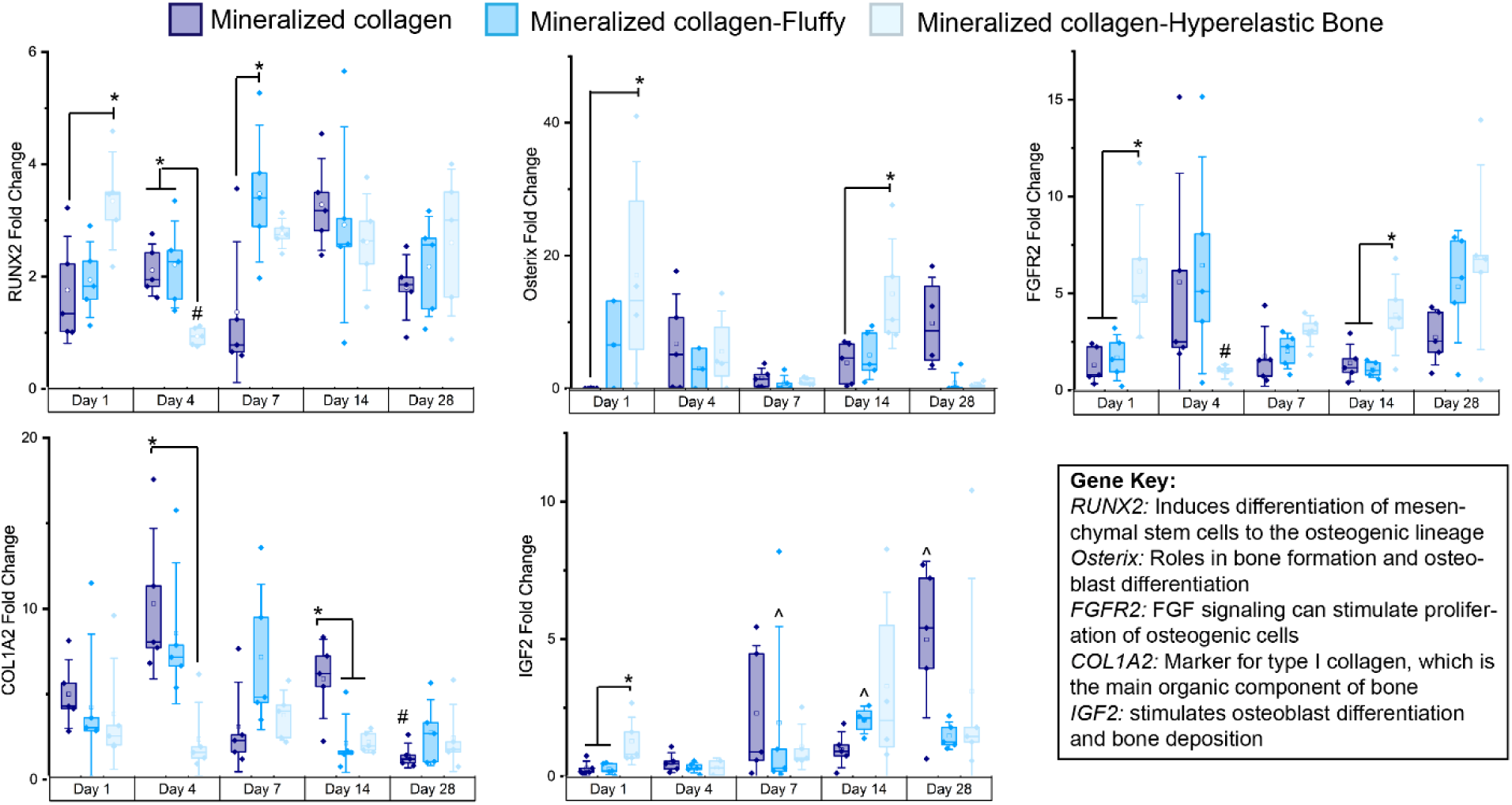
Osteogenic gene expression of mineralized collagen scaffolds (MC), Fluffy-PLG reinforced composites (MC-Fluffy), and Hyperelastic Bone reinforced composites (MC-HB) composites. Gene expression was determined by running RT-PCR on reverse transcribed RNA isolated from scaffolds and composites seeded with 100,000 hMSCs at days 1-28. Fold change of gene expression was normalized to 100,000 cells before seeding on scaffolds and composites, with a value of 1 representing the expression of 100,000 hMSCs. Values underneath this grey bar indicate downregulated genes. * indicates group(s) significant (p < 0.05) to another group on same day. # indicates a group on one day is significantly (p < 0.05) less than the same group on day 1. ^ indicates a group on one day is significantly (p < 0.05) greater than the same group on day 1. Error bars represent average ± standard deviation (n=5).

### 3.7. Fluffy-PLG reinforced composites formed the greatest amount of new calcium by the end of the study

The mineral content (calcium and phosphorous) of all hMSC-seeded constructs was evaluated at day 28, with results normalized to the values for acellular scaffolds or composites (**Fig. 4B**). MC-Fluffy composites displayed significantly (p < 0.05) greater calcium content than both MC scaffolds and MC-HB composites. Additionally, MC-Fluffy composites also contained significantly (p < 0.05) more phosphorous content than MC-HB composites. Although new mineral found in MC and MC-Fluffy samples was higher than MC-HB samples, MC-HB samples had overall much greater amounts of calcium and phosphorous than the other groups (**Supp. Table 9**).

## 4. Discussion

Advancing regenerative medicine technologies for CMF bone defects is challenging not only due to the large quantity of bone missing, but their irregular size and shape. There is an increasing need to identify biomaterial technologies to improve integration with the surrounding defect margins to aid cell recruitment and vascular ingrowth. Recent effort has begun to exploit shape-fitting technologies such as porous polymers that can reshape with changes in temperature [39-41]. Mineralized collagen scaffolds developed by our group as well as others have been shown to promote osteogenic processes and bone remodeling [29, 31, 32, 64-69]. However, the porous nature of these scaffolds that is important to aid cell activity results in poor bulk mechanical properties; further, these scaffolds lack inherent design features to promote conformal fitting with the defect margins [42]. Here, we examined inclusion of mechanical and biological reinforcement to a mineralized collagen scaffold via inclusion of 3D-printed structures formed using one of two variations of 3D-printed biomaterials, Fluffy-PLG and Hyperelastic Bone (90wt% CaP). We report compositional, mechanical, and biological performance of these scaffold composites, focusing on the direct role of the 3D-printed structures on mechanical properties but also the potential active role the reinforcing structure could have on an osteogenic response via the effect of degradation byproducts on cell activity.

As both Hyperelastic Bone and Fluffy-PLG contain the same PLG chemistry, and roughly equivalent amounts of PLG polymer per unit volume [46, 47, 51], we first investigated the degradation byproducts of Hyperelastic Bone to determine if there would be any impact on the metabolic activity of our osteogenic mineralized collagen scaffolds. Over the course of an accelerated 28-day degradation experiment, we observed significant elution of lactic and glycolic acid, as well as short term decrease in pH, though not significantly different than the change in solution pH observed for mineralized collagen scaffolds in the same conditions. More importantly, we observed no negative effect of the elution byproducts, across a range of dilutions, on hMSC metabolic activity. These effects were consistent regardless of the use of an accelerate (90°C) or convention (37°C) degradation conditions. These findings are largely consistent with previous studies using printed structures formed from Hyperelastic Bone 3D-prints that showed significant new bone formation *in vivo* [47, 51]. As a result, we did not expect Hyperelastic Bone or Fluffy-PLG 3D-prints to negatively impact osteogenesis and cell viability of mineralized collagen scaffolds. It is interesting to note that we also observed significant release of calcium and phosphorous content from the Hyperelastic Bone structures, though not as much calcium release as from the native mineralized collagen scaffold, which may be due to the more stable nature of the mineral component in Hyperelastic Bone versus that in the mineralized collagen scaffold.

Successful clinical use of a craniofacial bone regeneration scaffold requires the biomaterial to be surgically practical, notably readily customized to fit complex 3D defects and strong enough to withstand physiological loading [15, 37]. To address this challenge, we have developed an innovative composite approach, embedding a mechanically-robust polymer (e.g., PCL, PLA) 3D-printed reinforcement with millimeter-scale porosity into the mineralized collagen scaffold [42, 45]. The resultant collagen-3D-print composite displays the osteogenic activity of the scaffold, the strength of the 3D-print, and promotes regenerative healing in a porcine bone defect [28, 44, 45]. However, poor conformal contact significantly inhibits cell recruitment, angiogenesis, regenerative healing and greatly increases the risk of graft resorption [19, 38]. Common surgical interventions such as resorbable protective plates do not improve microscale conformal contact [70]. We showed selectively removing fibers from a PLA 3D-print creates variants with compressive strength in the longitudinal axis, but serial compression-expansion capacity in the radial [45]. These 3D-prints can be radially compressed and inserted into cylindrical defects, springing back to achieve close conformal contact. As a result, here we examined the mechanical performance of a new class of composite formed from the mineralized collagen scaffold and Fluffy-PLG or Hyperelastic Bone 3D-prints.

Hyperelastic Bone provided the greatest reinforcement to mineralized collagen scaffolds, most likely due to the reinforcements dominating the compressive mechanics similar to previous work with poly(lactic acid) composites [45]. MC-HB composites also displayed the highest loads achieved during a push-out test; however, the MC-HB composites demonstrated a more brittle nature, likely due to the form factor and not material. We expect that reducing the spacing between layers could reduce the brittleness and avoid many stress-concentrating regions created with wider spacing. While MC-Fluffy composites were significantly more flexible, they displayed no additional push-out strength compared to the mineralized scaffold itself. We also found resulting mechanical properties to be largely insensitive to changes in the mesh spacing (2 mm vs. 3 mm) over the range tested for the Fluffy-PLG prints, most likely due to their soft and flexible nature. Although MC-HB composites showed significantly improved mechanical performance, they did not achieve a moduli close to cancellous or cortical bone (0.1-2 GPa, 15-20 GPa [71]). While the need for mechanical properties of implant to match those of a target tissue may be relevant for inert materials which never remodel such as permanent joint replacements, design needs are very different for regenerative biomaterials that rely on cell mechanotransduction and remodeling. Indeed, it is essential to consider the multiscale properties of porous biomaterials. Porosity is essential for cell ingrowth, biotransport, and cell-mediated remodeling. Yet the mechanical performance of a porous material scales as the relative density (density of the porous material divided by the solid material from which it is fabricated) squared compared to the mechanical properties of the solid material that forms the individual pores and with which cells interact [72, 73]. Thus, a porous scaffold with cell-scale properties tuned to enhance MSC proliferation, remodeling, and matrix biosynthesis may present initial macroscale mechanical properties a factor of 1000 (or more) lower. The design of the mineralized scaffolds and polymer reinforcement is therefore tuned to improve surgical practicality and cell activity. The mineralized collagen scaffold used in this study has previously been shown to regenerate rabbit cranial defects without growth factor and cell supplementation, achieving directed healing >60% of the density and 50-80% the stiffness of native cranial bone within 3 months after implantation [33]. Similarly, Hyperelastic Bone has also been shown to promote long-term stability and bone formation in primate full-thickness cranial defects [48, 51]. With these results, we expect that the combination of two materials previously successfully tested separately *in vivo* would be able to promote greater bone regeneration together, and significant future efforts will be needed to identify the appropriate mesh structure to aid strength and conformal fitting capacity. Future efforts will concentrate on alterations to the polymer composition and mineral content to further boost endogenous OPG secretion to accelerate regenerative healing. An exciting opportunity are meshes based on Voronoi foam (random pore) architectures which exhibit well defined mechanical performance characteristics [74, 75]. Under load, the fibers that define the pores of these materials bend elastically during the first ∼10% of applied strain, then buckle plastically [74], suggesting implementing these designs with the 3D-Paints may be useful for conformal fitting capabilities.

This work showed exciting osteogenic potential of hMSC-seeded MC-HB and MC-Fluffy composites, maintained in culture in the absence of conventional osteogenic supplements (exogenous BMP2, osteogenic media). Both composites promoted a higher metabolic activity compared to native mineralized collagen scaffolds in early timepoints of the study (days 1-14); while by day 28 there were no differences between groups, all groups demonstrated significantly (∼3-fold) increased metabolic activity compared to the start of the experiment. We also quantified gene expression profiles for a series of osteogenic-linked genes as a function of biomaterial environment. *RUNX2* is a major transcription factor for bone and guides the differentiation of hMSCs to osteoblasts [76, 77]; we saw some evidence of increases in *RUNX2* expression at some early time points in reinforced composites. Downstream of *RUNX2, Osterix* regulates mature osteoblast differentiation and is thus connected to bone formation [78]. MC-HB composites upregulated *Osterix* at day 1 and 14, possibly indicating greater mature osteoblast differentiation in these scaffolds, however, all groups were upregulated at day 28. In addition to this, MC-HB composites upregulated Fibroblast Growth Factor Receptor-2 (*FGFR2*) at day 1 and 14, which is important due to FGF being a key regulator of bone development and FGF-2 coated scaffolds being shown to stimulate osteogenesis *in vivo* [79, 80]. *COL1A2* is a gene expression marker for type I collagen, the major collagen of bone [81]. While greater in MC scaffolds at days 4 and 14, all groups were upregulated throughout the study. Overall, we observed signatures of biomaterial induced increases in osteogenic signaling, though a MC-HB composite may have the greatest promise to accelerate early osteogenic activity. These findings are largely consistent with previous studies from our group showing the base mineralized scaffold itself is sufficient to promote hMSC osteogenic activity in the absence of osteogenic stimuli [82-85]. At minimum, we show inclusion of Fluffy-PLG or Hyperelastic Bone reinforcing structures into the scaffold does not reduce this response.

We also examined endogenous production of two proteins (OPG, VEGF) by hMSCs within the scaffolds and composites. OPG is a soluble glycoprotein and endogenous inhibitor of osteoclastogenesis and osteoclast activity [35, 86]. We have previously shown that MSC-osteoprogenitors in the mineralized collagen scaffold endogenously produce osteoprotegerin sufficient to inhibit osteoclast activity without negatively impacting MSC-osteogenesis [36]. Excitingly, we observed significant OPG production in all variants, though MC-HB composites promoted significantly increased OPG secretion over the first two weeks of culture. Calcium ion signaling has previously been suggested as a powerful signal to improve secretion of OPG by osteoblasts [87]. Our findings are consistent with this observation, suggesting increased OPG secretion by hMSCs in MC-HB composites may be due to increased release of Calcium and Phosphate ions during degradation (vs. MC scaffolds or MC-Fluffy composites). This indicates that inclusion of Hyperelastic Bone 3D-prints could not only regenerate bone through increased stiffness, which has been linked with greater mechanotransduction-induced bone formation in mineralized collagen scaffolds [88], but also provide additional inorganic ions to safely elevate OPG levels without the need of gene therapy or growth factors [56]. The timing of increased OPG production during significant Hyperelastic Bone degradation further suggests that the Hyperelastic Bone reinforcement structure may play an active role in promoting an osteogenic response in addition to passive mechanical reinforcement. We also explored VEGF production, as VEGF is a potential regulator of angiogenesis, and can also contribute to recruitment and activity of osteoblasts [89, 90]. Exogenous addition of VEGF can also improve bone formation [79]. While we observed significant increases in VEGF production over the 28-day experiment, the effect was insensitive to the inclusion of Fluffy-PLG or Hyperelastic Bone reinforcement structure.

Analysis of mineral content of the scaffold and composites after *in vitro* culture suggest significant new mineral formation in all constructs, though the greatest new absolute mineral formation was observed in MC-Fluffy composites. MC-HB composites had the least amount of new mineral formed compared to MC scaffolds and MC-Fluffy composites. However, the MC-HB composites had the greatest average amount of Calcium and Phosphorous present at day 28 before normalizing to unseeded controls (approx. 30% Ca and 15% P, 11% Ca and 5% P, 14% Ca and 7% P for MC-HB, MC-Fluffy, and MC, respectively), indicating that there is still a great amount of mineral present to induce osteogenesis. To further understand the low mineral formation at the end of the study in MC-HB composites, we plan to investigate the localization of calcium and phosphorous within the composite to determine if mineral is forming within the scaffold.

Together, our results demonstrate hMSC viability, gene expression, and protein expression that support the use of Hyperelastic Bone and Fluffy-PLG reinforced mineralized collagen composites for craniofacial bone repair applications. Of particular note, was the observation that MC-HB composites demonstrated increased MSC-osteoprogenitor secreted OPG and added stiffness. Ongoing efforts will more fully compare hMSC osteogenesis *in vitro* as well as *in vivo* bone regeneration for these disparate composites. Future efforts will also examine changes in the topology of the 3D-printed structure in order to add shape-fitting through structural means, similar to our recently published work with poly(lactic acid) composites [45]. This includes expanding the use of the 3D-print as a bioactive stimulus to increase endogenous OPG production as a means to transiently inhibit osteoclast activity and accelerate craniofacial bone repair.

## 5. Conclusions

We examined inclusion of reinforcing 3D-prints generated from two distinct 3D-paints (Hyperelastic Bone; Fluffy-PLG) in a mineralized collagen scaffold for craniofacial bone repair applications. Composites formed from Hyperelastic Bone or Fluffy-PLG structures were shown to offer passive and potentially active advantages to aid osteogenic activity. Notably, inclusion of a Hyperelastic Bone 3D-printed mesh significantly increased composite modulus and push-out force, though the brittle nature of the structure limited the conformal fitting capacity. Degradation byproducts of Hyperelastic Bone did not significantly reduce hMSC activity. Interesting, all composites and the native mineralized collagen scaffold supported significant osteogenic activity in the form of hMSC metabolic activity increases, shifts in osteogenic gene expression, and synthesis of new mineral. Further, while the scaffold and composites all promoted increased endogenous production and secretion of OPG and VEGF, Hyperelastic Bone reinforced composites showed significantly increased early secretion of OPG, suggesting these composites may increase hMSC osteogenesis and locally inhibit osteoclast activity to accelerate bone regeneration by mineral ion release and increased stiffness without the addition of growth factors or gene therapy.

## Supporting information

Statement of Significance

## Acknowledgements

This work was supported by the Office of the Assistant Secretary of Defense for Health Affairs Broad Agency Announcement for Extramural Medical Research through the Award No. W81XWH-16-1-0566. Research reported in this publication was also supported by the National Institute of Dental and Craniofacial Research of the National Institutes of Health under Award Number R21 DE026582. We are also grateful for funds provided by the NSF Graduate Research Fellowship DGE-1144245 (MJD) to perform this research. The interpretations and conclusions presented are those of the authors and are not necessarily endorsed by the Department of Defense, NIH, or NSF.

The authors would like to acknowledge the University of Illinois Roy J. Carver Biotechnology Center, the School of Chemical Science Microanalysis Lab, the Carl R. Woese Insitute for Genomic Biology, the Chemical and Biomolecular Engineering Department, and the Beckman Institute for Advanced Science and Technology, all located at the University of Illinois at Urbana-Champaign. The authors would also like to thank members of the Harley Lab who assisted in statistical advice and Matlab programs, Aidan, Samantha, and Aleczandria.

## Data Availability

The raw and processed data required to reproduce these findings are available to download from Dewey, Marley (2020), “Data Repository: Inclusion of a polylactide-co-glycolide co-polymer mesh enhances osteogenesis in mineralized collagen scaffolds”, Mendeley Data, V1, doi: 10.17632/tfx7dt4kyt.1.

## Disclosure

The authors Ramille N. Shah and Adam E. Jakus are co-founders and hold financial interests in Dimension Inx, LLC, which may be considered a potential competing interest.

## Notes

### Competing Interest Statement

RNS and AEJ are co-founders and hold financial interests in Dimension Inx, LLC.

### Summary of Updates

The title has been changed and more details were added to a few methods and the discussion.

## References

[1] T.A. Lew, J.A. Walker, J.C. Wenke, L.H. Blackbourne, R.G. Hale, Characterization of craniomaxillofacial battle injuries sustained by United States service members in the current conflicts of Iraq and Afghanistan, J Oral Maxillofac Surg 68(1) (2010) 3–7.

[2] A.J. Fong, B.T. Lemelman, S. Lam, G.M. Kleiber, R.R. Reid, L.J. Gottlieb, Reconstructive approach to hostile cranioplasty: A review of the University of Chicago experience, J Plast Reconstr Aesthet Surg 68(8) (2015) 1036–43.

[3] J.C. Lee, G.M. Kleiber, A.T. Pelletier, R.R. Reid, L.J. Gottlieb, Autologous immediate cranioplasty with vascularized bone in high-risk composite cranial defects, Plast Reconstr Surg 132(4) (2013) 967–75.

[4] A.H. Chao, P. Yu, R.J. Skoracki, F. Demonte, M.M. Hanasono, Microsurgical reconstruction of composite scalp and calvarial defects in patients with cancer: A 10-year experience., Head Neck (2012).

[5] E.I. Lee, A.H. Chao, R.J. Skoracki, P. Yu, F. DeMonte, M.M. Hanasono, Outcomes of calvarial reconstruction in cancer patients, Plast Reconstr Surg 133(3) (2014) 675–82.

[6] P. Tessier, Autogenous bone grafts taken from the calvarium for facial and cranial applications, Clin Plast Surg 9(4) (1982) 531–8.

[7] E.C. Kruijt Spanjer, G.K.P. Bittermann, I.E.M. van Hooijdonk, A.J.W.P. Rosenberg, D. Gawlitta, Taking the endochondral route to craniomaxillofacial bone regeneration: A logical approach?, Journal of Cranio-Maxillofacial Surgery 45 (2017) 1099–1106.

[8] A. Norozy, M.H.K. Motamedi, A. Ebrahimi, H. Khoshmohabat, Maxillofacial Fracture Patterns in Military Casualties, Journal of Oral and Maxillofacial Surgery 78(4) (2020) 611.e1-611.e6.

[9] B.D. Owens, J.F. Kragh, Jr., J.C. Wenke, J. Macaitis, C.E. Wade, J.B. Holcomb, Combat wounds in operation Iraqi Freedom and operation Enduring Freedom, J Trauma 64(2) (2008) 295–9.

[10] P.R. Brown Baer, J.C. Wenke, S.J. Thomas, C.R. Hale, Investigation of severe craniomaxillofacial battle injuries sustained by u.s. Service members: a case series, Craniomaxillofac Trauma Reconstr 5(4) (2012) 243–52.

[11] H. Fodstad, J.A. Love, J. Ekstedt, H. Fridén, B. Liliequist, Effect of cranioplasty on cerebrospinal fluid hydrodynamics in patients with the syndrome of the trephined., Acta Neurochir (Wien) 70(1-2) (1984) 21–30.

[12] M. Hagan, J.P. Bradley, Syndrome of the Trephined: Functional Improvement After Reconstruction of Large Cranial Vault Defects, J Craniofac Surg 28(5) (2017) 1129–1130.

[13] P.A. Zuk, Tissue engineering craniofacial defects with adult stem cells? Are we ready yet?, Pediatr Res 63(5) (2008) 478–86.

[14] M. Elsalanty, D. Genecov, Bone Grafts in Craniofacial Surgery, Craniomaxillofacial Trauma and Reconstruction 2 (2009) 125–134.

[15] M.H. Smith, C.L. Flanagan, J.M. Kemppainen, J.A. Sack, H. Chung, S. Das, S.J. Hollister, S.E. Feinberg, Computed tomography-based tissue-engineered scaffolds in craniomaxillofacial surgery, Int J Med Robot 3(3) (2007) 207–16.

[16] T.B. Dodson, R.A. Bays, R.C. Pfeffle, D.L. Barrow, Cranial bone graft to reconstruct the mandibular condyle in Macaca mulatta, J Oral Maxillofac Surg 55(3) (1997) 260–7.

[17] J.C. Banwart, M.A. Asher, R.S. Hassanein, Iliac crest bone graft harvest donor site morbidity. A statistical evaluation, Spine (Phila Pa 1976) 20(9) (1995) 1055–60.

[18] A. Depeyre, S. Touzet-Roumazeille, L. Lauwers, G. Raoul, J. Ferri, Retrospective evaluation of 211 patients with maxillofacial reconstruction using parietal bone graft for implants insertion, Journal of Cranio-Maxillofacial Surgery 44 (2016) 1162–1169.

[19] D. Zhang, O.J. George, K.M. Petersen, A.C. Jimenez-Vergara, M.S. Hahn, M.A. Grunlan, A bioactive “self-fitting” shape memory polymer scaffold with potential to treat cranio-maxillo facial bone defects, Acta Biomaterialia 10(11) (2014) 4597–4605.

[20] J.M. Piitulainen, T. Kauko, K.M.J. Aitasalo, V. Vuorinen, P.K. Vallittu, J.P. Posti, Outcomes of cranioplasty with synthetic materials and autologous bone grafts, World Neurosurg. 83(5) (2015) 708–714.

[21] S. Ghanaati, M. Barbeck, P. Booms, J. Lorenz, C.J. Kirkpatrick, R.A. Sader, Potential lack of “standardized” processing techniques for production of allogeneic and xenogeneic bone blocks for application in humans, Acta Biomaterialia 10 (2014) 3557–3562.

[22] K. Nelson, T. Fretwurst, A. Stricker, T. Steinberg, M. Wein, A. Spanou, Comparison of four different allogeneic bone grafts for alveolar ridge reconstruction: a preliminary histologic and biochemical analysis, Oral Surgery, Oral Medicine, Oral Pathology and Oral Radiology 118 (2014) 424–431.

[23] J.A. Fearon, D. Griner, K. Ditthakasem, M. Herbert, Autogenous Bone Reconstruction of Large Secondary Skull Defects, Plast Reconstr Surg 139(2) (2017) 427–438.

[24] K.S. Blum, S.J. Schneider, A.D. Rosenthal, Methyl methacrylate cranioplasty in children: long-term results., Pediatr Neurosurg 26(1) (1997) 33–5.

[25] A. Afifi, R.S. Djohan, W. Hammert, F.A. Papay, A.E. Barnett, J.E. Zins, Lessons learned reconstructing complex scalp defects using free flaps and a cranioplasty in one stage, J Craniofac Surg 21(4) (2010) 1205–9.

[26] R. Dimitriou, G.C. Babis, Biomaterial osseointegration enhancement with biophysical stimulation, Journal of Musculoskeletal Neuronal Interactions 7 (2007) 253–265.

[27] B.A. Harley, A.K. Lynn, Z. Wissner-Gross, W. Bonfield, I.V. Yannas, L.J. Gibson, Design of a multiphase osteochondral scaffold. II. Fabrication of a mineralized collagen-glycosaminoglycan scaffold, Journal of Biomedical Materials Research - Part A 92 (2010) 1066–1077.

[28] D.W. Weisgerber, S.R. Caliari, B.A.C. Harley, Mineralized collagen scaffolds induce hMSC osteogenesis and matrix remodeling, Biomaterials Science 3 (2015) 533–542.

[29] G. Cunniffe, G. Dickson, S. Partap, K. Stanton, F.J. O’Brien, Development and characterisation of a collagen nano-hydroxyapatite composite scaffold for bone tissue engineering, J Mater Sci Mater Med 21 (2010) 2293–8.

[30] J.P. Gleeson, T. Weber, T. Levingstone, A.A. Al-Munajjed, F.J. O’Brien, J. Hammer, N.A. Plunkett, C. Jungreuthmayer, Development of a biomimetic collagen-hydroxyapatite scaffold for bone tissue engineering using a SBF immersion technique, Journal of Biomedical Materials Research Part B: Applied Biomaterials 90B (2009) 584–591.

[31] F.G. Lyons, J.P. Gleeson, S. Partap, K. Coghlan, F.J. O’Brien, Novel microhydroxyapatite particles in a collagen scaffold: a bioactive bone void filler?, Clinical Orthopaedics and Related Research 472 (2014) 1318–28.

[32] X. Ren, D.W. Weisgerber, D. Bischoff, M.S. Lewis, R.R. Reid, T.C. He, D.T. Yamaguchi, T.A. Miller, B.A.C. Harley, J.C. Lee, Nanoparticulate Mineralized Collagen Scaffolds and BMP-9 Induce a Long-Term Bone Cartilage Construct in Human Mesenchymal Stem Cells, Advanced Healthcare Materials 5 (2016) 1821–1830.

[33] X. Ren, V. Tu, D. Bischoff, D. Weisgerber, M. Lewis, D. Yamaguchi, T. Miller, B. Harley, J. Lee, Nanoparticulate mineralized collagen scaffolds induce in vivo bone regeneration independent of progenitor cell loading or exogenous growth factor stimulation., Biomaterials 89 (2016) 67–78.

[34] C. Lo Sicco, D. Reverberi, C. Balbi, V. Ulivi, E. Principi, L. Pascucci, P. Becherini, M.C. Bosco, L. Varesio, C. Franzin, M. Pozzobon, R. Cancedda, R. Tasso, Mesenchymal Stem Cell-Derived Extracellular Vesicles as Mediators of Anti-Inflammatory Effects: Endorsement of Macrophage Polarization, Stem cells translational medicine 6(3) (2017) 1018–1028.

[35] X. Ren, Q. Zhou, D. Foulad, M.J. Dewey, D. Bischoff, T.A. Miller, D.T. Yamaguchi, B.A.C. Harley, J.C. Lee, Nanoparticulate mineralized collagen glycosaminoglycan materials directly and indirectly inhibit osteoclastogenesis and osteoclast activation, J Tissue Eng Regen Med 13(5) (2019) 823–34.

[36] X. Ren, Q. Zhou, D. Foulad, A.S. Tiffany, M.J. Dewey, D. Bischoff, T.A. Miller, R.R. Reid, T.-C. He, D.T. Yamaguchi, B.A.C. Harley, J.C. Lee, Osteoprotegerin reduces osteoclast resorption activity without affecting osteogenesis on nanoparticulate mineralized collagen scaffolds, Sci Adv 5(6) (2019) eaaw4991.

[37] S.J. Hollister, W.L. Murphy, Scaffold translation: barriers between concept and clinic, Tissue Eng Part B Rev 17(6) (2011) 459–74.

[38] B.T. Smith, J. Shum, M. Wong, A.G. Mikos, S. Young, Bone tissue engineering challenges in oral & maxillofacial surgery, in: L.E. Bertassoni, P.G. Coelho (Eds.), Engineering Mineralized and Load Bearing Tissues, Springer International Publishing, Cham, 2015, pp. 57–78.

[39] L.N. Nail, D. Zhang, J.L. Reinhard, M.A. Grunlan, Fabrication of a Bioactive, PCL-based “Self-fitting” Shape Memory Polymer Scaffold, Journal of Visualized Experiments 104 (2015) e52981.

[40] R. Xie, J. Hu, O. Hoffmann, Y. Zhang, F. Ng, T. Qin, X. Guo, Self-fitting shape memory polymer foam inducing bone regeneration: A rabbit femoral defect study, Biochimica et Biophysica Acta (BBA)-General Subjects 1862(4) (2018) 936–945.

[41] D. Zhang, O.J. George, K.M. Petersen, A.C. Jimenez-Vergara, M.S. Hahn, M.A. Grunlan, A bioactive “self-fitting” shape memory polymer scaffold with potential to treat cranio-maxillo facial bone defects, Acta Biomaterialia 10 (2014) 4597–4605.

[42] D.W. Weisgerber, K. Erning, C. Flanagan, S.J. Hollister, B.A.C. Harley, Evaluation of multi-scale mineralized collagen-polycaprolactone composites for bone tissue engineering, J Mech Behav Biomed Mater 61 (2016) 318–327.

[43] D. Weisgerber, D. Milner, H. Lopez-Lake, M. Rubessa, S. Lotti, K. Polkoff, R. Hortensius, C. Flanagan, S. Hollister, M. Wheeler, B. Harley, A mineralized collagen-polycaprolactone composite promotes healing of a porcine mandibular ramus defect, Tissue Eng Part A 0 (2017) 1–12.

[44] D.W. Weisgerber, K. Erning, C.L. Flanagan, S.J. Hollister, B.A.C. Harley, Evaluation of multi-scale mineralized collagen-polycaprolactone composites for bone tissue engineering, Journal of the Mechanical Behavior of Biomedical Materials 61 (2016) 318–327.

[45] M.J. Dewey, E.M. Johnson, D.W. Weisgerber, M.B. Wheeler, B.A.C. Harley, Shape-fitting collagen-PLA composite promotes osteogenic differentiation of porcine adipose stem cells, Journal of the Mechanical Behavior of Biomedical Materials 95 (2019) 21–33.

[46] A.E. Jakus, N.R. Geisendorfer, P.L. Lewis, R.N. Shah, 3D-printing porosity: A new approach to creating elevated porosity materials and structures, Acta Biomaterialia 72 (2018) 94–109.

[47] R. Alluri, A. Jakus, S. Bougioukli, W. Pannell, O. Sugiyama, A. Tang, R. Shah, J.R. Lieberman, 3D printed hyperelastic “bone” scaffolds and regional gene therapy: A novel approach to bone healing, J Biomed Mater Res Part A 106A (2018) 1104–1110.

[48] A.E. Jakus, A.L. Rutz, S.W. Jordan, A. Kannan, S.M. Mitchell, C. Yun, K.D. Koube, S.C. Yoo, H.E. Whiteley, C.-P. Richter, R.D. Galiano, W.K. Hsu, S.R. Stock, E.L. Hsu, R.N. Shah, Hyperelastic “bone”: A highly versatile, growth factor–free, osteoregenerative, scalable, and surgically friendly biomaterial, Science Translational Medicine 8(358) (2016) 358ra127.

[49] X. Liu, A.E. Jakus, M. Kural, H. Qian, A. Engler, M. Ghaedi, R. Shah, D.M. Steinbacher, L.E. Niklason, Vascularization of Natural and Synthetic Bone Scaffolds, Cell Transplant 27(8) (2018) 1269–1280.

[50] J.A. Driscoll, R. Lubbe, A.E. Jakus, K. Chang, M. Haleem, C. Yun, G. Singh, A.D. Schneider, K.M. Katchko, C. Soriano, M. Newton, T. Maerz, X. Li, K. Baker, W.K. Hsu, R.N. Shah, S.R. Stock, E.L. Hsu, 3D-Printed Ceramic-Demineralized Bone Matrix Hyperelastic Bone Composite Scaffolds for Spinal Fusion, Tissue Eng Part A 26(3-4) (2020) 157–166.

[51] Yu-Hui Huang, A.E. Jakus, S.W. Jordan, Z. Dumanian, K. Parker, L. Zhao, P.K. Patel, R.N. Shah, Three-Dimensionally Printed Hyperelastic Bone Scaffolds Accelerate Bone Regeneration in Critical-Size Calvarial Bone Defects, American Society of Plastic Surgeons (2019) 1397–1407.

[52] A.S. F1635-04a, Standard Test Method for in vitro Degradation Testing of Hydrolytically Degradable Polymer Resins and Fabricated Forms for Surgical Implants, ASTM International, West Conshohocken, PA, 2018, pp. 1–5.

[53] L.N. Borshchevskaya, T.L. Gordeeva, A.N. Kalinina, S.P. Sineokii, Spectrophotometric Determination of Lactic Acid, Zhurnal Analitticheskoi Khimii 71 (2016) 787–790.

[54] J. Viccaro, E. Ambye, Colorimetric Determination of Glycolic Acid with B-Naphthol, Microchemical Journal 17 (1972) 710–718.

[55] X. Ren, Q. Zhou, D. Foulad, M.J. Dewey, D. Bischoff, T.A. Miller, D.T. Yamaguchi, B.A.C. Harley, J.C. Lee, Nanoparticulate mineralized collagen glycosaminoglycan materials directly and indirectly inhibit osteoclastogenesis and osteoclast activation, Journal of tissue engineering and regenerative medicine 13(5) (2019) 823–834.

[56] X. Ren, Q. Zhou, D. Foulad, A.S. Tiffany, M.J. Dewey, D. Bischoff, T.A. Miller, R.R. Reid, T.-c. He, D.T. Yamaguchi, B.A.C. Harley, J.C. Lee, Osteoprotegerin reduces osteoclast resorption activity without affecting osteogenesis on nanoparticulate mineralized collagen scaffolds, Science Advances 5 (2019) 1–12.

[57] H.J. Conrad, W.-J. Seong, J.S. Hodges, S. Grami, S.C. Jeong, Comparison of Push-In versus Pull-Out Tests on Bone-Implant Interfaces of Rabbit Tibia Dental Implant Healing Model, Clinical Implant Dentistry and Related Research 15 (2011) 460–469.

[58] A.S. Tiffany, D.L. Gray, T.J. Woods, K. Subedi, B.A.C. Harley, The inclusion of zinc into mineralized collagen scaffolds for craniofacial bone repair applications, Acta Biomaterialia 93 (2019) 86–96.

[59] M.J. Dewey, E.M. Johnson, S.T. Slater, D.J. Milner, M.B. Wheeler, B.A.C. Harley, Mineralized collagen scaffolds fabricated with amniotic membrane matrix increase osteogenesis under inflammatory conditions, Regenerative Biomaterials rbaa005 (2020) 1–12.

[60] M.J. Dewey, A.V. Nosatov, K. Subedi, B. Harley, Anisotropic mineralized collagen scaffolds accelerate osteogenic response in a glycosaminoglycan-dependent fashion, RSC Advances 10(26) (2020) 15629–15641.

[61] ISO 10993-5, Part 5: Tests for in vitro cytotoxicity, Biological evaluation of medical devices, International Organization for Standardization, Switzerland, 2009, pp. 1–42.

[62] T.F. Scientific, Isolation of Total RNA from Difficult Tissues. https://www.thermofisher.com/us/en/home/references/ambion-tech-support/rna-isolation/tech-notes/isolation-of-total-rna-from-difficult-tissues.html. (Accessed March 30 2020).

[63] R.L. Ott, M.T. Longnecker, An Introduction to Statistical Methods and Data Analysis, 7 ed., Cengage Learning 2016.

[64] A. Al-Munajjed, J. Gleeson, F. O’Brien, Development of a collagen calcium-phosphate scaffold as a novel bone graft substitute., Stud Health Technol Inform 133 (2008) 11–20.

[65] D. Florent, T. Levingstone, W. Schneeweiss, M. de Swartw, H. Jahns, J. Gleeson, F. O’Brien, Enhanced bone healing using collagen– hydroxyapatite scaffold implantation in the treatment of a large multiloculated mandibular aneurysmal bone cyst in a thoroughbred filly, Journal of tissue engineering and regenerative medicine 9 (2015) 1193–1199.

[66] B. Hoyer, A. Bernhardt, S. Heinemann, I. Stachel, M. Meyer, M. Gelinsky, Biomimetically mineralized salmon collagen scaffolds for application in bone tissue engineering., Biomacromolecules 13 (2012) 1059–1066.

[67] F.J. O’Brien, E. Thompson, S.A. Cryan, A. López-Noriega, E. Quinlan, H.M. Kelly, Development of collagen–hydroxyapatite scaffolds incorporating PLGA and alginate microparticles for the controlled delivery of rhBMP-2 for bone tissue engineering, Journal of Controlled Release 198 (2014) 71–79.

[68] X. Ren, D. Bischoff, D. Weisgerber, V. Tu, M. Lewis, D. Yamaguchi, T. Miller, B.A.C. Harley, J. Lee, Osteogenesis on nanoparticulate mineralized collagen scaffolds via autogenous activation of the canonical BMP receptor signaling pathway, Biomaterials 50 (2015) 107–14.

[69] D. Weisgerber, S. Caliari, B. Harley, Mineralized collagen scaffolds induce hMSC osteogenesis and matrix remodeling, Biomaterials Science 3 (2015) 533–42.

[70] J.H. Phillips, B.A. Rahn, Fixation effects on membranous and endochondral onlay bone graft revascularization and bone deposition., Plast Reconstr Surg 85(6) (1990) 891–7.

[71] S. Bose, M. Roy, A. Bandyopadhyay, Recent advances in bone tissue engineering scaffolds, Trends Biotechnol. 30 (2012) 546–554.

[72] L.A. Gibson, M. & Harley, B., Cellular Materials in Nature and Medicine, 1 ed., Cambridge University Press 2010.

[73] B.A. Harley, J.H. Leung, E.C.C.M. Silva, L.J. Gibson, Mechanical characterization of collagen-glycosaminoglycan scaffolds, Acta Biomaterialia 3 (2007) 463–474.

[74] L.J. Gibson, M.F. Ashby, B.A. Harley, Cellular materials in nature and medicine, Cambridge University Press, Cambridge, U.K., 2010.

[75] L.J. Gibson, M.F. Ashby, Cellular solids: structure and properties, 2nd ed., Cambridge University Press, Cambridge, U.K., 1997.

[76] L.D. Carbonare, G. Innamorati, M.T. Valenti, Transcription Factor Runx2 and its Application to Bone Tissue Engineering, Stem Cell Reviews and Reports 8 (2012) 891–897.

[77] A.W. James, Review of Signaling Pathways Governing MSC Osteogenic and Adipogenic Differentiation, Scientifica 2013 (2013) 1–17.

[78] B. Yao, J. Wang, S. Qu, Y. Liu, Y. Jin, J. Lu, Q. Bao, L. Li, H. Yuan, C. Ma, Upregulated osterix promotes invasion and bone metastasis and predicts for a poor prognosis in breast cancer, Cell Death Dis 10(1) (2019) 28.

[79] Y.-R. Yun, J.H. Jang, E. Jeon, W. Kang, S. Lee, J.-E. Won, H.W. Kim, I. Wall, Administration of growth factors for bone regeneration, Regenerative Medicine 7 (2012).

[80] G. Draenert, K. Draenert, T. Tischer, Dose-dependent osteoinductive effects of bFGF in rabbits, Growth Factors 27 (2009) 419–24.

[81] U. Khetarpal, C. Morton, COL1A2 and COL2A1 Expression in Temporal Bone of Lethal Osteogenesis Imperfecta, Arch Otolaryngol Head Neck Surg 119 (1993) 1305–1314.

[82] D.W. Weisgerber, S.R. Caliari, B.A.C. Harley, Mineralized collagen scaffolds induce hMSC osteogenesis and matrix remodeling, Biomater Sci 3(3) (2015) 533–42.

[83] S.R. Caliari, B.A.C. Harley, Structural and biochemical modification of a collagen scaffold to selectively enhance MSC tenogenic, chondrogenic, and osteogenic differentiation, Advanced healthcare materials 3(7) (2014) 1086–96.

[84] S.R. Caliari, B.A.C. Harley, Collagen-GAG scaffold biophysical properties bias MSC lineage selection in the presence of mixed soluble signals, Tissue Eng A 20(17-18) (2014) 2463–72.

[85] J.M. Banks, L.C. Mozdzen, B.A.C. Harley, R.C. Bailey, The combined effects of matrix stiffness and growth factor immobilization on the bioactivity and differentiation capabilities of adipose-derived stem cells, Biomaterials 35(32) (2014) 8951–9.

[86] T. Standal, C. Seidel, Ø. Hjertner, T. Plesner, R.D. Sanderson, A. Waage, M. Borset, A. Sundan, Osteoprotegerin is bound, internalized, and degraded by multiple myeloma cells, BLOOD 100 (2002) 3002–3007.

[87] J.J. Bergh, Y. Xu, M.C. Farach-Carson, Osteoprotegerin expression and secretion are regulated by calcium influx through the L-type voltage-sensitive calcium channel, Endocrinology 145(1) (2004) 426–36.

[88] Q. Zhou, S. Lyu, A. Bertrand, A. Hu, C. Chan, X. Ren, M. Dewey, A. Tiffany, B. Harley, J. Lee, Stiffness of Nanoparticulate Mineralized Collagen Scaffolds Triggers Osteogenesis via Mechanotransduction and Canonical Wnt Signaling, BioRxiv (2020).

[89] K. Hu, B.R. Olsen, The roles of vascular endothelial growth factor in bone repair and regeneration, Bone 91 (2016) 30–38.

[90] Y.-Q. Yang, Y.-Y. Tan, R. Wong, A. Wenden, L.-K. Zhang, A.B.M. Rabie, The role of vascular endothelial growth factor in ossification, Int J Oral Sci 4(2) (2012) 64–68.

