## Supplementary material for "Inclusion of a 3D-printed Hyperelastic bone mesh improves mechanical and osteogenic performance of a mineralized collagen scaffold": Statement of Significance

Craniofacial bone defects occur due to trauma, congenital abnormalities, and during the course of surgical treatments for stroke and cancer. Clinically-available technologies for craniofacial reconstruction are non-regenerative. We report inclusion of polymer mesh generated via 3D-printing into a mineralized collagen scaffold under development for craniofacial bone regeneration. Scaffold-mesh composites improved mechanical performance and mesenchymal stem cell activity, notably secretion of osteoprotegerin, a soluble glycoprotein and endogenous inhibitor of osteoclast activity. These findings suggest the exciting possibility to co-optimize the composition and architecture of an integrated polymer mesh to both passively aid surgical-practicality and actively accelerate regenerative healing.
